# Development of Neonatal Functional Brain Network Organisation and Its Association with Childhood Neurodevelopmental Outcomes

**DOI:** 10.64898/2026.10.20.760497

**Authors:** Qianwen Chang, Sunniva Fenn-Moltu, Andrew Chew, Alexandra F Bonthrone, Barat Gal-Er, Chiara Nosarti, Tomoki Arichi, Grainne McAlonan, Serena J. Counsell, Dafnis Batalle

**Author notes:** **Corresponding author:** Dr Dafnis Batalle, Department of Forensic and Neurodevelopmental Science, Institute of Psychiatry, Psychology & Neuroscience, King’s College London 16, De Crespigny Park, SE5 8AF, London, United Kingdom, **Email:**.

## Abstract

The human brain undergoes rapid dynamic changes in the neonatal period that are essential for later cognitive and behavioural performance. Using resting-state fMRI data from 394 neonates (325 term-born, 69 preterm-born) within the first month after birth, we characterised the early development of functional network organisation, the effects of preterm birth, and the association between neonatal functional organisation and later neurodevelopmental outcomes in infancy (18 months) and childhood (4-6 years). We found increased functional integration and segregation across a wide range of functional sub-networks and increased mean functional connectivity strength (mean FCS) within the subcortical and cerebellar regions with increasing PMA at scan. Compared with term-born neonates, preterm-born neonates showed significantly lower mean FCS, functional integration and segregation globally. We further showed that whilst neonatal functional brain network organisation was not associated with neurodevelopmental outcomes measured during infancy (18 months), it was associated with later neurodevelopmental outcomes in early childhood (4-6 years). Specifically, higher functional integration and segregation at birth were linked with lower SDQ total difficulties score at 4-6 years. Our findings suggest that neonatal functional brain organisation reflects not only transient developmental processes, but also stable individual differences that provide insight into neurodevelopmental outcomes in later childhood.

## 1 Introduction

The perinatal period is a critical time for the formation of the developing brain’s neural networks through a programmed sequence of rapid and dynamic changes that are essential for later cognitive and behavioural performance (Gilmore et al., 2018). Functional brain networks emerge before birth but remain largely immature and continue to develop across the first years after birth (Cao et al., 2017; Eyre et al., 2021; Gao, 2025; Gilmore et al., 2018; Zhao et al., 2019). Functional segregation and integration are key principles of functional brain organisation: densely connected subnetworks specialise for specific functions, and these distributed subnetworks coordinate to support complex cognitive functions (Sporns, 2013). During early development, there is a strengthened balance between integration and segregation (Cao et al., 2017; Gao et al., 2011; Zhao et al., 2019), with the balance between these two main descriptors of topological organisation thought to be fundamental for supporting emerging cognitive functions (Cohen & D’Esposito, 2016; Wang et al., 2021). Neonatal functional brain organisation is sensitive to perinatal factors such as preterm birth, and has been shown in numerous studies to link to alterations in functional connectivity and network topology at term-equivalent age (Eyre et al., 2021; Fenn-Moltu et al., 2023; Sa de Almeida et al., 2021; Smyser et al., 2010; Vanes et al., 2023) which persist even into adulthood (Karolis et al., 2016; Schafer et al., 2009). Atypical network organisation has also been associated with an increased likelihood of neurodevelopmental conditions and poor cognitive outcomes (Michael et al., 2024; Rudie et al., 2013; Weber et al., 2024).

Several studies have reported significant associations between neonatal functional brain organisation and later neurodevelopmental outcomes at 1.5-2 years of age. Functional integration and segregation (Li et al., 2024) and other network topology features (e.g., participation coefficient; Fang et al., 2025) have been linked to cognitive and/or language scores measured by the Bayley Scales of Infant and Toddler Development, Third Edition (Bayley-III, Bayley, 2006). However, these studies have reported relatively small effect size or modest prediction accuracy, or have identified associations involving regional, rather than whole-brain topology features; other studies report weak or no associations (e.g., Fenn-Moltu et al., 2023). The relationship between overall neonatal functional network organisation and later neurodevelopmental outcomes therefore remains largely unknown. One potential explanation is that neurodevelopmental assessments in toddlerhood are inherently limited because many cognitive abilities emerge later in development and are therefore not captured at this stage (Gao, 2025; Gustavson et al., 2025), or perhaps because specific network features or regional features are more predictive than others. It is also not clear whether specific aspects of neonatal functional brain network organisation reflect transient developmental processes or more stable individual differences that are linked with later neurodevelopmental outcomes. Furthermore, it has not been established whether neonatal functional brain network organisation associates with neurodevelopment into early childhood, when cognitive functions and behavioural abilities are more established.

In this study, we aimed to: (i) characterise the development of neonatal functional brain organisation; (ii) examine the effects of preterm birth on neonatal functional brain network organisation; and (iii) explore associations with later neurodevelopmental outcomes in both toddlerhood (18 months) and early childhood (4-6 years). Using resting-state fMRI data from the developing Human Connectome Project (dHCP; Edwards et al., 2022), we calculated functional integration and segregation in the neonatal period, examined their developmental trajectories and the effects of preterm birth at both whole-brain and sub-network levels, and their associations with later neurodevelopmental outcomes. We further examined outcomes in both toddlerhood and early childhood to understand whether assessments at 4-6 years of age may capture individual differences in cognitive and behavioural performance more reliably than assessments at 18 months.

## 2 Methods

### 2.1 Participants

A total of 394 neonates (325 term-born, 69 preterm-born) from the developing Human Connectome Project (dHCP) cohort (Edwards et al., 2022) were included in this study. Inclusion and exclusion criteria are provided in Figure 1. Resting-state fMRI (rs-fMRI) was acquired at term equivalent age (postmenstrual age 37-45 weeks). Demographic information and follow-up assessments are shown in Table 1.

**Figure 1.**
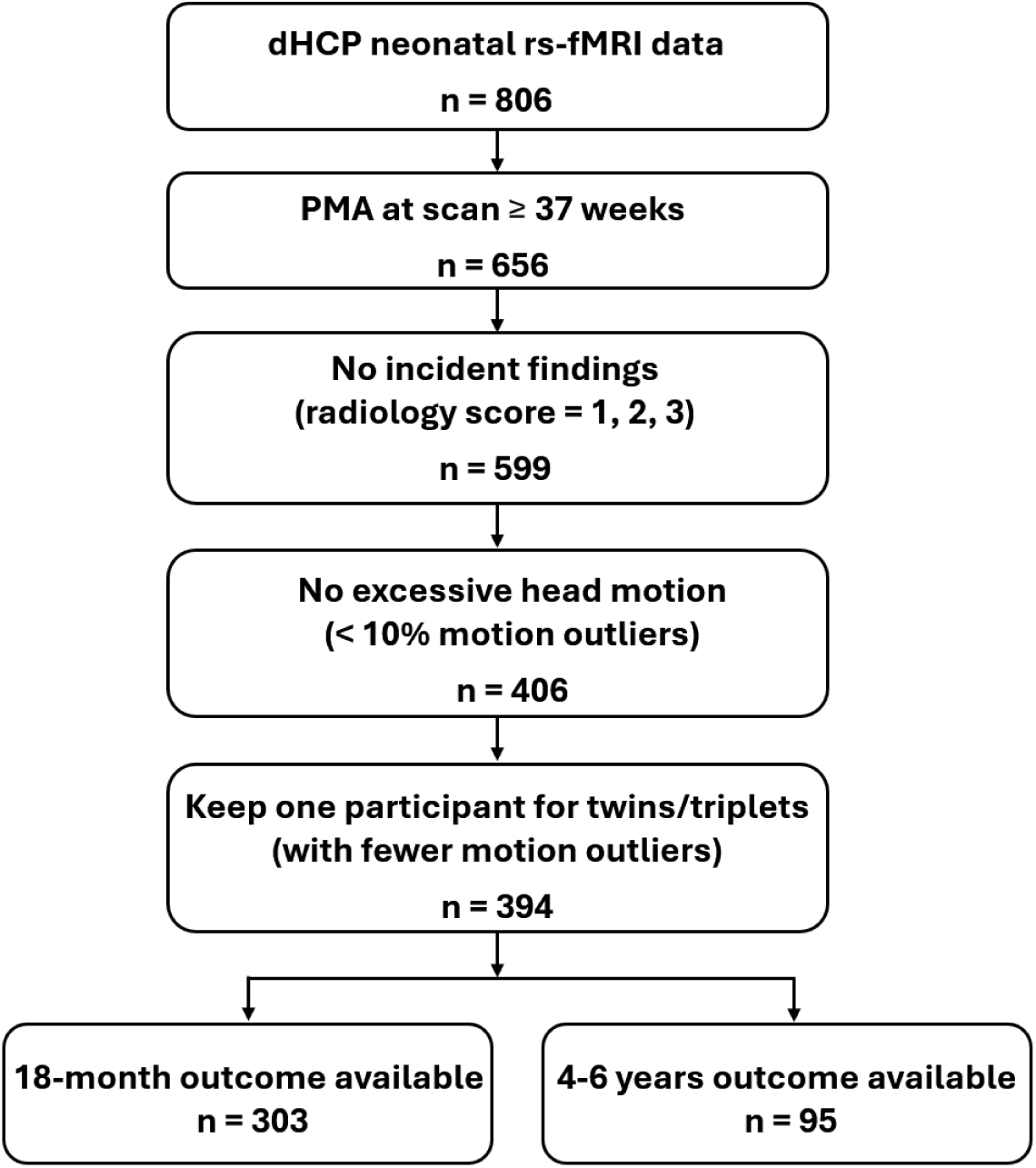
**Inclusion and exclusion criteria.**

**Table 1.** Demographic information and follow-up assessment at 18 months and 4-6 years of age.

|  | Term<br>(n = 325) | Preterm<br>(n = 69) | Statistic | p-value |
| --- | --- | --- | --- | --- |
| GA at birth [weeks] | 40.04 (1.26) | 32.77 (3.30) | 13.100 | < 0.001*** |
| PMA at scan [weeks] | 41.60 (1.67) | 40.61 (2.11) | 3.640 | < 0.001*** |
| Sex (number of males, %) | 174 (54.00%) | 40 (58.00%) | 0.290 | 0.590 |
| Index of Multiple Deprivation (IMD) | 13657.4 (7403.2) | 15723.5 (8138.5) | -2.020 | 0.044* |
| % framewise displacement [FD] outliers | 5.37 (2.62) | 4.61 (2.55) | 2.210 | 0.027* |
| **18-month assessment<sup>+</sup>** |  |  |  |  |
| Corrected age at assessment [months] | 19.06 (2.16) | 19.25 (3.30) | 0.453 | 0.651 |
| Uncorrected age at assessment [months] | 19.05 (2.18) | 20.97 (3.45) | -7.180 | < 0.001*** |
| Bayley III cognition | 101.68 (10.70) | 100.21 (11.89) | 0.844 | 0.399 |
| Bayley language | 99.74 (15.43) | 96.00 (14.17) | 1.490 | 0.138 |
| Bayley motor | 102.45 (9.86) | 100.66 (9.87) | 1.350 | 0.179 |
| Q-CHAT | 29.80 (8.40) | 30.49 (10.27) | -0.259 | 0.796 |
| CBCL externalising | 11.42 (6.78) | 10.15 (7.21) | 1.360 | 0.174 |
| CBCL internalising | 5.58 (4.91) | 5.38 (5.14) | 0.497 | 0.620 |
| CBCL other behavioural problems | 7.88 (4.83) | 7.83 (5.04) | 0.044 | 0.965 |
| CBCL sleep score | 2.67 (2.02) | 2.89 (2.92) | 0.199 | 0.843 |
| **4-6 years assessment<sup>++</sup>** |  |  |  |  |
| Age at assessment [years] | 4.68 (0.74) | 4.68 (0.70) | 0.112 | 0.915 |
| CBQ surgency | 53.07 (10.74) | 54.86 (9.06) | -0.657 | 0.514 |
| CBQ negative affectivity | 46.70 (10.78) | 45.05 (11.36) | 0.605 | 0.548 |
| CBQ effortful control | 62.68 (8.66) | 65.75 (6.14) | -1.110 | 0.268 |
| SCQ total score | 4.86 (3.18) | 6.21 (3.71) | -1.570 | 0.117 |
| EmQue emotional contagion | 1.63 (1.55) | 2.05 (2.66) | -0.014 | 0.992 |
| EmQue attention to others' feelings | 9.33 (2.28) | 9.26 (1.97) | 0.202 | 0.844 |
| EmQue prosocial actions | 6.40 (2.69) | 6.63 (2.03) | -0.201 | 0.844 |
| SDQ total difficulties score | 7.84 (4.72) | 7.16 (4.09) | 0.551 | 0.585 |
| ADHD total score | 10.66 (8.02) | 11.35 (6.18) | -0.745 | 0.459 |
| BRIEF-P inhibition | 24.12 (5.26) | 24.37 (4.91) | -0.093 | 0.929 |
| BRIEF-P shifting | 14.04 (3.92) | 14.68 (4.76) | -0.614 | 0.542 |
| BRIEF-P emotional control | 15.95 (4.15) | 16.95 (5.03) | -0.776 | 0.441 |
| BRIEF-P working memory | 23.46 (5.39) | 24.74 (5.01) | -1.260 | 0.208 |
| BRIEF-P planning/organising | 15.01 (2.95) | 15.84 (3.37) | -0.731 | 0.468 |

**Note:** values are presented as mean (SD) if not indicated otherwise.
Chi-square test for sex, Mann-Whitney U test for other variables, \* $p < 0.05$ , \*\* $p < 0.01$ , \*\*\* $p < 0.001$ .
<sup>+</sup>Assessments at 18 months were available for 303 subjects (256 term-born, 47 preterm-born).
<sup>++</sup>Assessments at 4-6 years were available for 95 subjects (76 term-born, 19 preterm-born).

Complete neurodevelopmental assessments at 18 months were available for 303 subjects (Table 1). This included: the Bayley Scales of Infant and Toddler Development, Third Edition (Bayley-III, Bayley, 2006) to assess general developmental outcomes across motor, language, and cognitive domains, with higher scores indicating greater skills; the Child Behaviour Checklist (CBCL, Achenbach & Ruffle, 2000) to evaluate emotional and behavioural problems, including internalising, externalising, sleep, and other problem subscales, with higher scores indicating a higher frequency of behavioural, emotional and social problems; and the Quantitative Checklist for Autism in Toddlers (Q-CHAT, Allison et al., 2008) to measure atypical social, sensory, and repetitive behaviours, with higher total scores indicating a greater frequency of autistic-related traits.

Follow-up assessments at 4-6 years were available for 95 subjects, which were acquired as controls for the CHiP (Congenital Heart Imaging Project) study (Chew et al., 2024, 2026). These measures were further categorised into socio-emotional and executive function domains. For socio-emotional domains, Children’s Behaviour Questionnaire-Very Short Form (CBQ-VSF, Putnam et al., 2014) measured children’s temperament across three subscales (surgency, negative affectivity, and effortful control). The ADHD Rating Scale (ADHD-RS, McGoey et al., 2007) was used to assess the frequency of ADHD symptoms based on DSM-IV (Diagnostic and Statistical Manual of Mental Disorders, Fourth Edition) criteria, providing a combined score from two subscales (inattention and hyperactivity/impulsivity). The Strengths and Difficulties Questionnaires (SDQ, Goodman, 1997, 2001) assessed emotional and behavioural problems across five subscales (conduct problems, hyperactivity/inattention, emotional symptoms, peer problems, and prosocial behaviour). A total difficulties score was then calculated by summing the first four subscales. The Empathy Questionnaire (EmQue, Rieffe et al., 2010) assessed empathy-related behaviours, summarised into three scales (emotion contagion, attention to others’ feelings, and prosocial actions). Executive functions were assessed using the Behaviour Rating Inventory of Executive Function - preschool edition (BRIEF-P, Goodman, 2001), which includes inhibition, shifting, emotional control, working memory, and planning.

To assess potential selection bias associated with follow-up data, we compared demographic information and in-scanner head motion between subjects with and without follow-up data at 18 months and with and without follow-up data at 4-6 years, both in the whole sample and within term-and preterm-born groups separately (Supplementary Tables S2-S7). Subjects with and without follow-up data were broadly comparable, although participants with follow-up at 18 months had a higher proportion of females (χ^2^(1) = 5.844, *p* = 0.016) and had a lower level of social deprivation, i.e., a higher postcode-based index of multiple deprivation (IMD) (Z = 2.820, *p* = 0.005). Participants with follow-up data at 4-6 years had higher PMA at scan (Z = 2.980, *p* = 0.003).

### 2.2 MRI data acquisition and preprocessing

MRI data were collected at the Evelina Newborn Imaging Centre, Evelina London Children’s Hospital, using a 3 Tesla Philips Achieva system (Philips Healthcare, Best NL) with a 32-channel neonatal head coil and imaging system (Rapid Biomedical GmbH, Rimpar DE) (Hughes et al., 2017). **Resting-state fMRI** was acquired using an echo planar imaging sequence with multiband excitation (factor 9) (repetition time (TR) = 392 ms, echo time (TE) = 38 ms, voxel size = 2.15 × 2.15 × 2.15 mm, flip angle = 34^°^, 45 slices, number of volumes = 2300). **T1-weighted images** had a reconstructed spatial resolution = 0.8 × 0.8 × 0.8 mm, field of view = 145 ×122 × 100 mm, TR = 4795 ms. **T2-weighted images** had a reconstructed spatial resolution = 0.8 × 0.8 × 0.8 mm, field of view = 145 × 145 × 108 mm, TR = 12 s, TE = 156 ms (Edwards et al., 2022).

fMRI data were preprocessed using the dHCP neonatal fMRI pipeline (Fitzgibbon et al., 2020), which included dynamic distortion and motion correction using slice-to-volume and rigid-body registration, high-pass filtering with a 150s cutoff and ICA denoising. The data were further bandpass filtered to 0.01-0.1Hz for functional connectivity analysis (Fenn-Moltu et al., 2023). Data were subsequently registered to T2w native space using boundary-based registration and then nonlinearly transformed into a 40-week neonatal template from the extended dHCP volumetric atlas (Schuh et al., 2018). fMRI data with excessive motion (more than 10% volumes classified as motion outliers, i.e., framewise displacement [FD] > 75th centile + 1.5 IQR) or incidental MRI findings of clinical significance (radiology score = 4 or 5) were excluded (Fenn-Moltu et al., 2023; França et al., 2024).

### 2.3 Data analysis

#### 2.3.1 Whole-brain atlas

To ensure generalisability, an anatomically defined whole-brain atlas was constructed. The atlas comprises 400 cortical regions from the Schaefer-400 atlas (Schaefer et al., 2018), as used in a number of other neonatal studies (King et al., 2023; Truzzi & Cusack, 2023; Yates et al., 2023; Yin et al., 2025). 12 Subcortical regions were taken from the dHCP template (Schuh et al., 2018), and 21 cerebellar regions (i.e., bilateral lobules I-IV, V, VI, Crus I, Crus II, VIIB, VIIIA, VIIIB, IX, X, and vermis) from the SUIT atlas (Diedrichsen, 2006), resulting in a total of 433 regions. The atlas was nonlinearly transformed to the extended dHCP 40-week template space (Schuh et al., 2018) using the Advanced Normalisation Tools (ANTs) SyN algorithm (Avants et al., 2011). Cortical parcels were then assigned by their spatial overlap to one of the Yeo-7 cortical sub-networks (Yeo et al., 2011). Subcortical and cerebellar parcels were treated as separate sub-networks, resulting in 9 sub-networks in total.

#### 2.3.2 Functional connectivity and functional network features

Functional connectivity was computed as the pairwise Pearson’s correlation of the average BOLD timeseries between the 433 brain parcels, with only positive correlations retained. This was done as the biological meaning of negative functional connectivity is unknown, and thus its interpretation remains controversial.

Functional network measures, including mean functional connectivity strength (mean FCS), normalised global efficiency, and normalised local efficiency, were calculated using the Brain Connectivity Toolbox (Rubinov & Sporns, 2010). Normalised weighted global and local efficiency were used to quantify functional integration and segregation. All network features were calculated as the averaged value across a range of network densities (ranging from 5% to 50%, with 1% increments). In order to calculate normalised features, functional connectivity matrices were normalised by total network strength at each network density (Batalle et al., 2017). To further examine the spatial effects of early brain development, mean FCS, and normalised global and local efficiency were calculated at both parcel and sub-network levels (by averaging over parcels within each of the Yeo-7 sub-networks) (Achard & Bullmore, 2007; Chen et al., 2019).

#### 2.3.3 Development of functional network features and effect of preterm birth

We further examined the effects of typical development and preterm birth status (term/preterm birth) on functional network features. For developmental changes, linear regression models were fitted in all term-born neonates: *Network features ∼ PMA at scan + Sex + in-scanner head motion (FD outliers) [in term-born neonates].* For the effect of preterm birth, linear regression models were fitted in the whole sample: *Network features ∼ PMA at scan + Birth status (term/preterm) + Sex + FD outliers [in the whole-sample].* For network- and parcel-level analyses, models were fit separately for each of the 9 sub-networks or 433 parcels. P-values were adjusted for multiple comparisons using the Benjamini-Hochberg FDR method (Benjamini & Hochberg, 1995) with each set of analyses.

#### 2.3.4 Associations between functional network features and neurodevelopmental outcomes at 18 months and 4-6 years

Associations between network features at birth and neurodevelopmental outcomes at 18 months and 4-6 years were assessed using Spearman’s rank partial correlations, controlling for corrected age at assessment, PMA at scan, number of FD outliers, sex, and IMD. Analyses were conducted separately for assessments at 18 months, socio-emotional measures at 4-6 years and executive functions at 4-6 years. Multiple comparison correction was performed using the Benjamini-Hochberg FDR method (Benjamini & Hochberg, 1995) with each set of analyses.

To examine whether associations between network features at birth and neurodevelopmental outcomes differed between term-and preterm-born groups, additional linear regression models included network features, preterm birth status, and their interaction, controlling for corrected age at assessment, PMA at scan, number of FD outliers, sex, and IMD.

## 3 Results

### 3.1 Typical development of functional networks and effects of preterm birth across the whole brain

Both normalised global (*r* = 0.219, *p* < 0.001) and local efficiency (*r* = 0.247, *p* < 0.001) increased with PMA at scan across the whole brain in term-born infants (Fig. 2). No significant association with age was found for the mean FCS (*r* = −0.07, *p* = 0.211). Preterm-born neonates showed significantly lower mean FCS (standardised *β* = −0.203, *p* < 0.001), normalised global (standardised *β* = −0.126, *p* = 0.011), and normalised local efficiency (standardised *β* = −0.144, *p* = 0.003) in comparison to term-born neonates.

**Figure 2.**
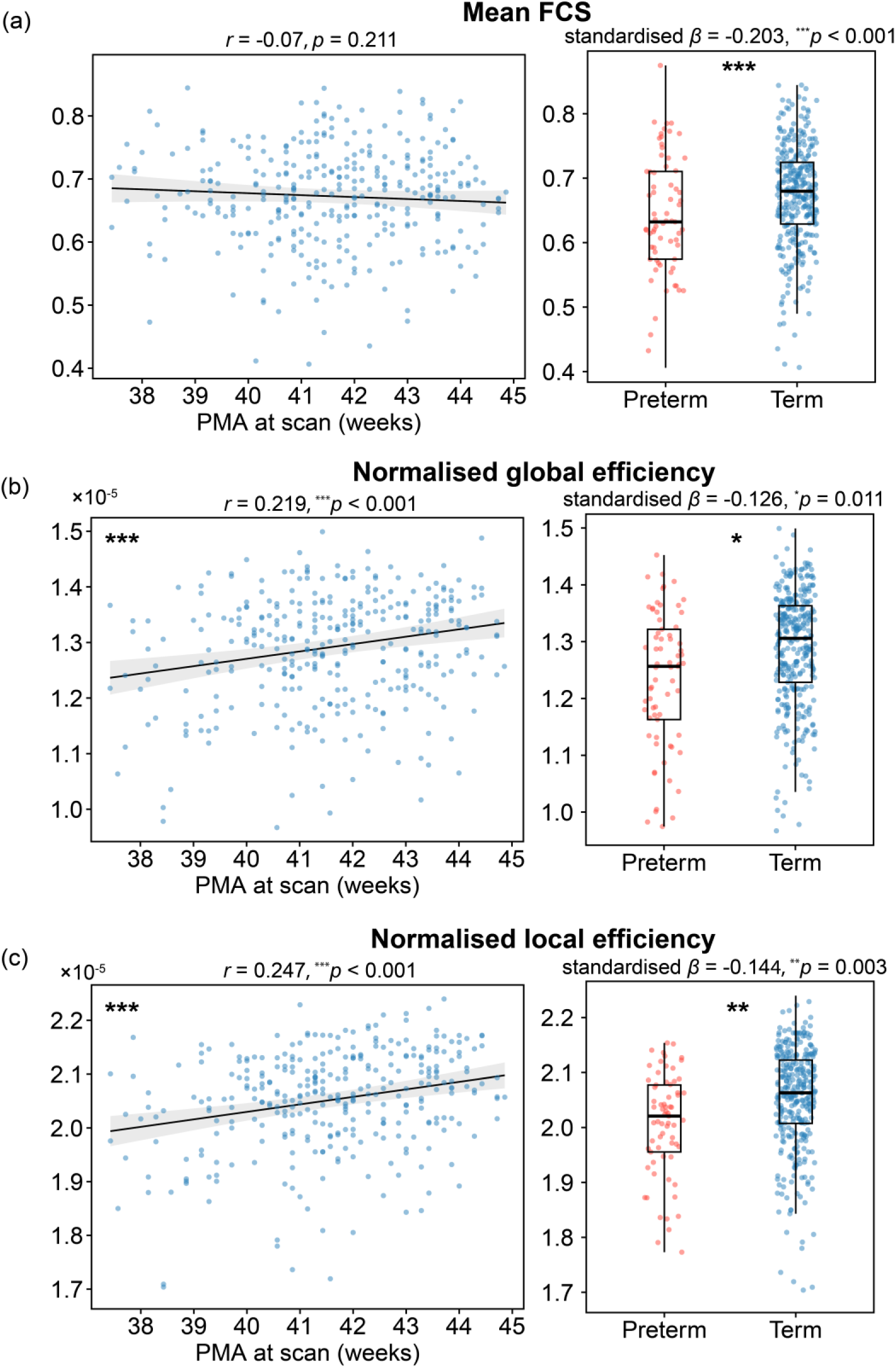
Effects of PMA at scan (left) and preterm birth (right) on functional network features at the whole-brain level. (a) Mean functional connectivity strength (mean FCS) (b) Normalised global efficiency (c) Normalised local efficiency. For the effect of PMA at scan, linear regression models were fitted in term-born neonates, controlling for sex and head motion (FD outliers). For effect of preterm birth, linear regression models were fitted in all neonates, controlling for PMA at scan, sex, and head motion. \**p*<0.05, \*\**p*<0.01, \*\*\**p*<0.001.

### 3.2 Typical development of functional networks and effects of preterm birth at the sub-network and parcel levels

Mean FCS in the subcortical (*r* = 0.159, *p*_FDR_ = 0.018) and cerebellar sub-networks (*r* = 0.348, *p*_FDR_ < 0.001) increased significantly with PMA at scan (Fig. 3a). Normalised global efficiency in the visual (VN; *r* = 0.244, *p*_FDR_ < 0.001), dorsal attention (DAN; *r* = 0.165, *p*_FDR_ = 0.005), frontoparietal (FPN; *r* = 0.168, *p*_FDR_ = 0.005), default mode (DMN; *r* = 0.128, *p*_FDR_ = 0.032), subcortical (SCN; *r* = 0.243, *p*_FDR_ < 0.001), and cerebellar networks (CBN; *r* = 0.368, *p*_FDR_ < 0.001) increased significantly with PMA at scan (*p*_FDR_ values < 0.05; Fig. 3b). Normalised local efficiency significantly increased in the VN (*r* = 0.219, *p*_FDR_ < 0.001), DAN (*r* = 0.131, *p*_FDR_ = 0.033), FPN (*r* = 0.134, *p*_FDR_ = 0.033), SCN (*r* = 0.218, *p*_FDR_ < 0.001) and CBN (*r* = 0.368, *p*_FDR_ < 0.001), while decreased in the salience/ventral attention sub-network (SN/VAN) with PMA at scan (*r* = −0.125, *p*_FDR_ = 0.037; Fig. 3c). At the parcel level, significant increases with PMA at scan were found in the cerebellar regions across all three metrics. For normalised global and local efficiency, additional positive clusters were observed in the visual sub-network and bilateral prefrontal cortex. In contrast, negative clusters were identified within the sensorimotor and salience/ventral attention sub-networks (Fig. 3).

**Figure 3.**
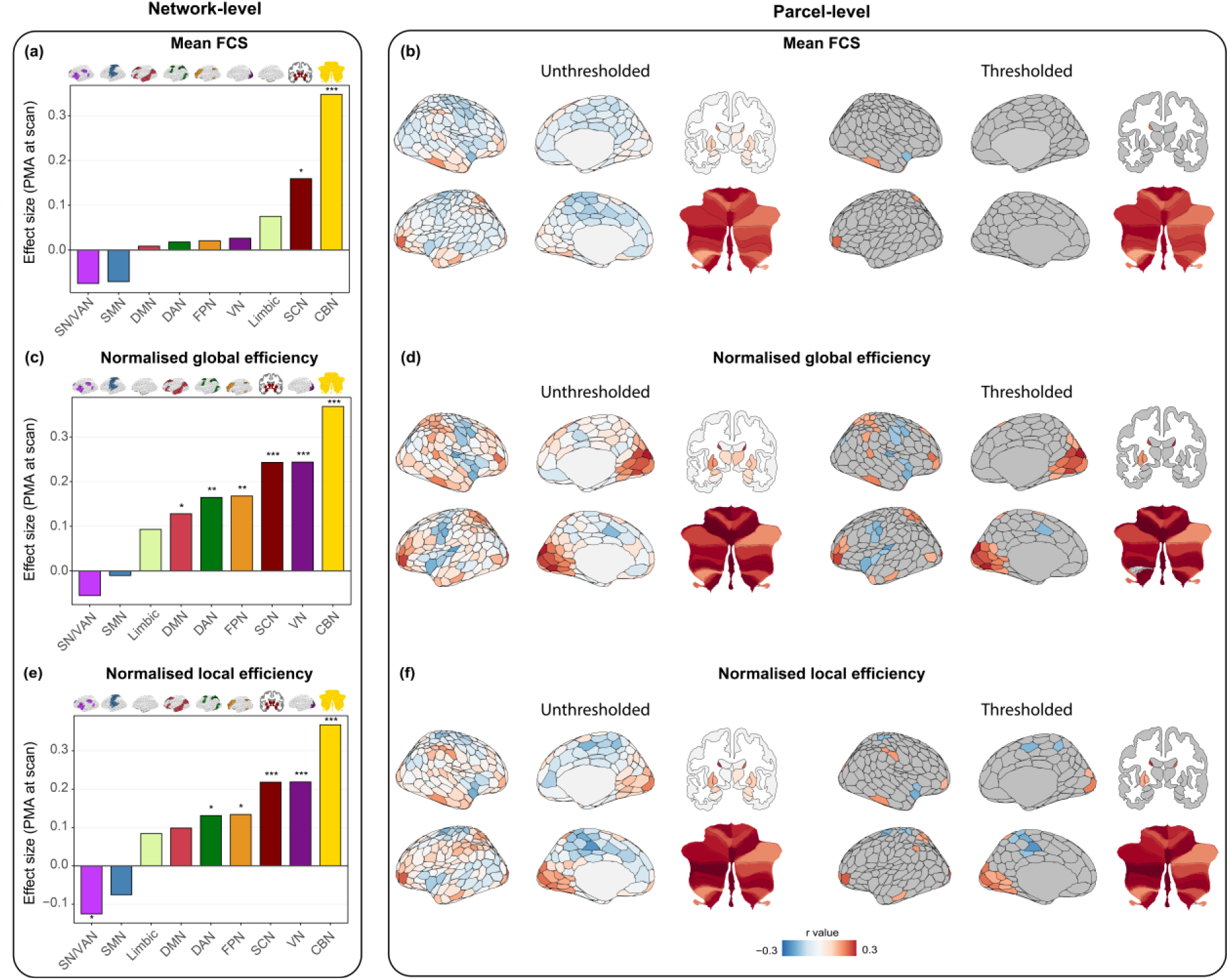
Effects of PMA at scan on functional network features at the sub-network (left) and parcel (right) levels. (a, b) Mean functional connectivity strength (mean FCS); (c, d) normalised global efficiency; and (e, f) normalised local efficiency. Linear regression models were fitted in term-born neonates, with PMA at scan as the main predictor, controlling for sex and in-scanner head motion (FD outliers). Effect sizes were reported as partial correlation coefficients (*r*). Bar plots showed the network-level results, and brain maps showed the parcel-level results. Sub-network level *p* values were corrected using the Benjamini-Hochberg FDR method across the 9 sub-networks. Unthresholded brain maps show results for all parcels, while thresholded brain maps show only parcels with *p*_FDR_ < 0.05, corrected for 433 parcels. SN/VAN = salience/ventral attention network; SMN = sensorimotor network; DMN = default mode network; DAN = dorsal attention network; FPN = frontoparietal network; VN = visual network; SCN = subcortical network; CBN = cerebellar network. \**p*_FDR_ < 0.05, \*\**p*_FDR_ < 0.01, \*\*\**p*_FDR_ < 0.001, FDR corrected for 9 sub-networks.

Compared with term-born neonates, preterm-born neonates showed significantly lower mean FCS across all the 9 sub-networks (*p*_FDR_ values < 0.001; Fig. 4a). Normalised global efficiency in the SN/VAN (standardised *β* = −0.239, *p*FDR < 0.001), DMN (standardised *β* = −0.133, *p*FDR = 0.023), and CBN (standardised *β* = −0.131, *p*FDR = 0.023) were significantly lower in preterm-born neonates (Fig. 4b). Normalised local efficiency in the SN/VAN reduced was significantly reduced in preterm-born neonates (standardised *β* = −0.219, *p*_FDR_ < 0.001; Fig. 4c). At the parcel level, preterm birth was associated with a widespread decrease in mean FCS across the whole brain. Preterm birth effect was restricted for normalised global and local efficiency at the parcel level. For normalised global efficiency, significant decreases in preterm-born neonates were found in the bilateral insula, the cingulate cortex, cerebellar Crus II and VIIb, as well as the left thalamus and lentiform. For normalised local efficiency, significant decreases in preterm-born neonates were only found in two nodes in the left insula (Fig. 4).

**Figure 4.**
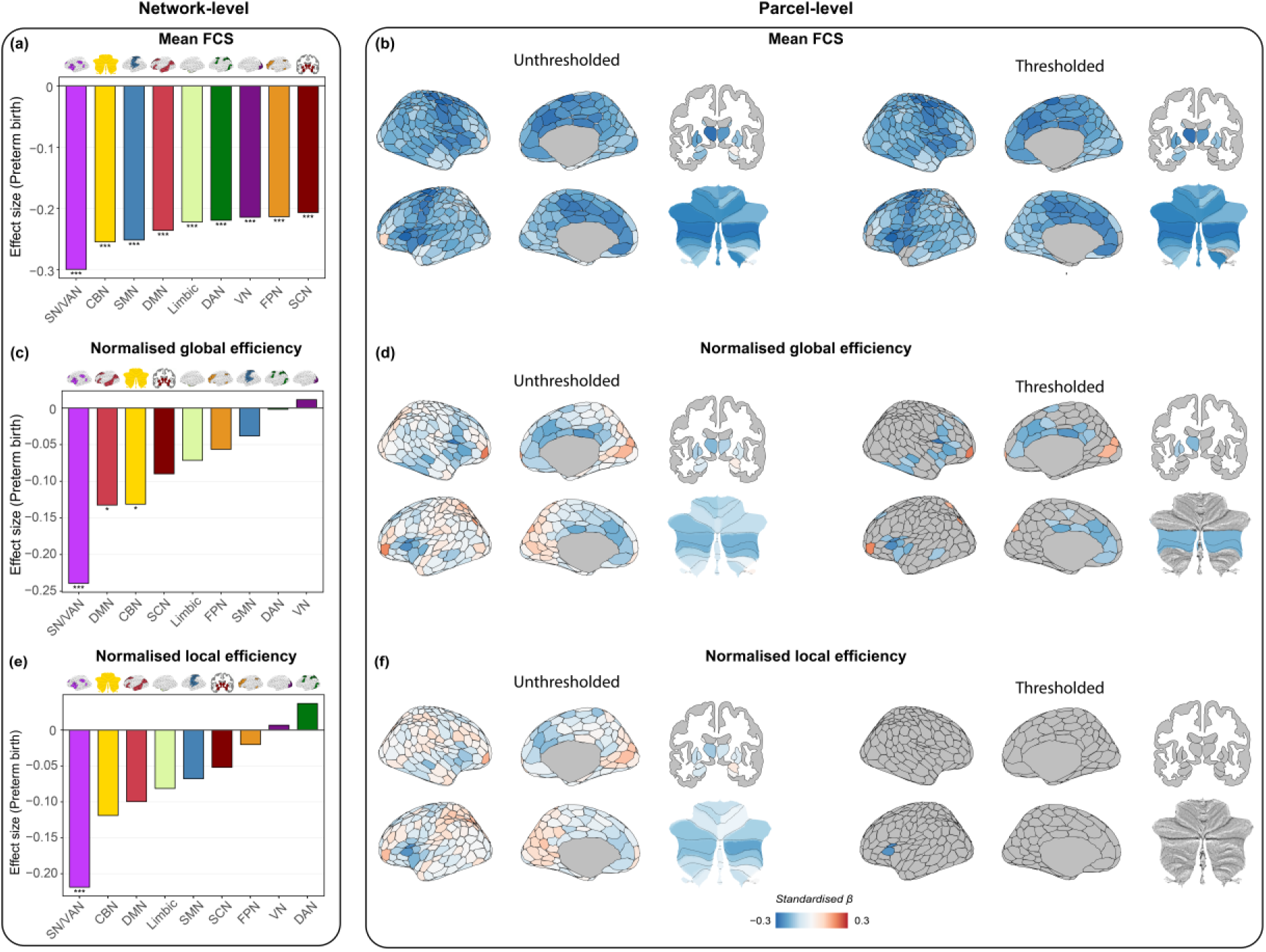
Effects of preterm birth on functional network features at the sub-network (left) and parcel (right) levels. (a, b) Mean functional connectivity strength (mean FCS); (c, d) normalised global efficiency; and (e, f) normalised local efficiency. Linear regression models were fitted in all neonates, with preterm birth status as the main predictor, controlling for PMA at scan, sex, and in-scanner head motion (FD outliers). Effect sizes were reported as standardised regression coefficient (*β*). Bar plots showed the sub-network level results, and brain maps showed parcel level results. Positive values indicate higher values in preterm-born neonates, while negative values indicate lower values compared with term-born neonates. Sub-network-level *p* values were corrected using the Benjamini-Hochberg FDR method across the 9 sub-networks. Unthresholded brain maps show results for all parcels, while thresholded brain maps show only parcels with *p*_FDR_ < 0.05, corrected for 433 parcels. SN/VAN = salience/ventral attention network; SMN = sensorimotor network; DMN = default mode network; DAN = dorsal attention network; FPN = frontoparietal network; VN = visual network; SCN = subcortical network; CBN = cerebellar network. \**p*_FDR_ < 0.05, \*\**p*_FDR_ < 0.01, \*\*\**p*_FDR_ < 0.001.

### 3.3 Associations between functional network features and neurodevelopmental outcomes at 18 months and 4-6 years of age

Normalised local efficiency (*r* = −0.306, *p*_FDR_ = 0.035) and local efficiency (*r* = −0.302, *p*_FDR_ = 0.035) were negatively correlated with SDQ total difficulties score at 4-6 years across both term- and preterm-born neonates (Fig. 5). Regional analysis further showed that normalised global efficiency in the frontoparietal sub-network was negatively associated with SDQ total difficulties score (*r* = −0.362, *p*_FDR_ = 0.006). No further significant associations were found either with executive functions at 4-6 years of age or with neurodevelopmental outcomes at 18 months of age. Detailed results for all associations, including non-significant findings, are provided in Supplementary Table S1.

**Figure 5.**
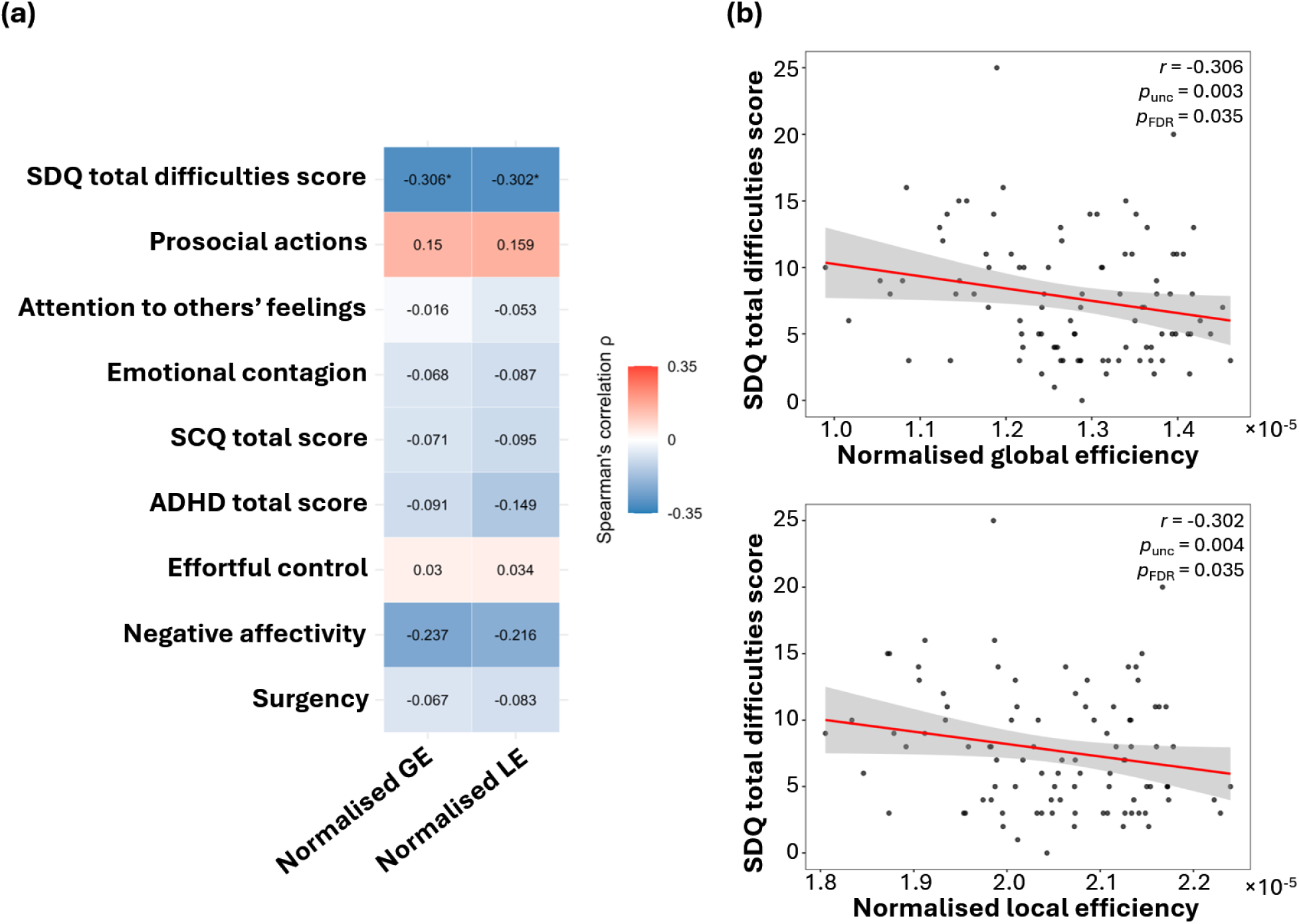
(a) Associations between functional network features at birth and neurodevelopmental outcomes at 4-6 years. Spearman’s partial correlations were calculated, controlling for corrected age at assessment, PMA at scan, head motion (FD outliers), sex, and IMD. \**p*_FDR_ < 0.05, FDR corrected. (b) Significant associations between SDQ total difficulties score and normalised global and local efficiency. *p*_unc_: uncorrected *p*-value; *p*_FDR_: FDR-corrected *p*-value.

Additional analyses showed no significant interactions between network features and preterm birth status for outcomes at 4-6 years of age (all *p*unc values > 0.1). After removing the interaction term, the main effect of preterm birth status was also non-significant (all *p*unc values > 0.05).

## 4 Discussion

In this study, we characterised the early development of the human brain’s functional network properties (i.e., mean functional connectivity strength, functional integration and functional segregation) within the first month after birth and examined the effects of preterm birth at the whole-brain and network level. We found that functional network integration and segregation globally increased with PMA at scan, but not mean FCS. In comparison with neonates born at full term, preterm-born neonates showed significantly lower global mean FCS, functional integration and segregation. We further showed that although neonatal functional organisation was not associated with neurodevelopmental outcomes measured during toddlerhood (18 months), it was significantly associated with later neurodevelopmental outcomes in early childhood (4-6 years). Specifically, higher functional integration and segregation at birth were linked with lower SDQ total difficulties score at 4-6 years. Our findings suggest that neonatal functional brain organisation reflects not only transient developmental processes but also provide insight into childhood neurodevelopmental outcomes.

We observed that mean functional connectivity strength increased significantly with PMA at scan within the subcortical and the cerebellar regions, although no significant changes were found globally. These findings align with previous studies demonstrating that functional connectivity between the cerebellum and higher-order networks is already present at birth and continues to strengthen during early development, supporting the emergence of cognitive functions (Lyu et al., 2025; Tikoo et al., 2025). In addition, thalamocortical connections undergo rapid development, suggesting that they may have a critical role in establishing cortical specialisation during the neonatal period (Oldham et al., 2025; Park et al., 2024; Toulmin et al., 2015).

Both global functional integration and segregation increased significantly with PMA at scan. At the network level, increases in functional integration were observed in 6 out of 9 sub-networks (visual, subcortical, cerebellar, dorsal attention, frontoparietal, and default mode sub-networks), while increases in functional segregation were found in 5 out of 9 sub-networks (visual, subcortical, cerebellar, dorsal attention, and frontoparietal sub-networks). The most marked increase in functional integration and segregation was seen within the visual network. This finding is consistent with broader evidence demonstrating that primary unimodal sensory and motor networks mature earlier during the neonatal period, providing essential input and output for higher-order functions and establishes the foundation for the development of higher-order association networks (Gao, 2025). Increased functional integration and segregation were also observed in higher-order association sub-networks, including the dorsal attention, frontoparietal, and default-mode networks, suggesting ongoing development of large-scale functional organisation in the period immediately after birth. Using precision mapping of individual-level resting-state networks, previous studies have also found pronounced postnatal development of the higher-order association networks (including the default-mode network) (Derman & Ferradal, 2025; Zhao et al., 2025). Of interest, normalised local efficiency decreased significantly with PMA at scan in the salience/ventral attention network. The salience/ventral attention network interacts with the dorsal attention, default-mode, and frontoparietal networks and may support attention and bottom-up salience detection during early development (Gao, 2025; Onofrj et al., 2022; Scheinost et al., 2024). While within-network functional connectivity of the salience network increases, functional connectivity between the salience network and both the default mode and frontoparietal networks gradually decreases across the first two years after birth (Gao et al., 2015; Scheinost et al., 2024), reflecting increased network specialisation. The decreased functional segregation observed in our study may reflect developmental topological changes that are not captured by functional connectivity strength alone. Given that the salience network remains relatively immature during the neonatal period, reduced local efficiency may reflect network reorganisation before later network specialisation. Preterm-born neonates showed significantly lower functional integration, functional segregation, and mean functional connectivity strength during the postnatal period. Previous studies using independent component analysis (ICA) to identify resting-state networks (RSNs) have reported widespread impairments in functional connectivity within both primary sensorimotor networks (including motor, somatosensory, auditory, and visual networks) and higher-order association regions (such as frontoparietal regions) in preterm-born neonates (Eyre et al., 2021; Fenn-Moltu et al., 2023). Using the pre-defined Yeo-7 anatomical atlas, we similarly observed reduced mean functional connectivity strength globally and across all subnetworks in preterm-born neonates. Consistent with prior work, preterm-born neonates showed lower functional integration and segregation compared with term-born neonates (Bouyssi-Kobar et al., 2019; López-Guerrero & Alcauter, 2025). These alterations may reflect disrupted developmental processes during the equivalent period to the third trimester, which is known to be a critical period for myelination, synaptogenesis, the establishment of long-range connectivity and thalamocortical fibres (Keunen et al., 2017). Importantly, these differences have long-term implications as the effects of preterm birth on network organisation have been reported to extend well beyond the postnatal period into childhood (Disselhoff et al., 2025), adolescence (Schafer et al., 2009) and adulthood (Hadaya et al., 2025).

Neonatal functional brain organisation was associated with childhood but not earlier toddler neurodevelopmental outcomes. In particular, higher functional integration and segregation were associated with lower SDQ total difficulties score at 4-6 years of age. The SDQ difficulties score captures both internalising (emotional and peer relationship problems) and externalising symptoms (conduct problems and hyperactivity), with higher scores suggesting greater psychopathology (Goodman & Goodman, 2009). At the sub-network level, the association was found within the frontoparietal network, a core system involved in cognitive control and emotion regulation that has been consistently implicated in internalising and externalising difficulties (Cole et al., 2014; Marek & Dosenbach, 2018; Xie et al., 2023). For example, Bashford-Largo et al. (2024) linked lower conduct problems measured by SDQ with greater grey matter volume in the frontotemporal regions in adolescents. Wee et al. (2017) found that the clustering coefficient of structural connectivity in the right inferior frontal gyrus during the neonatal period was associated with externalising behaviours at 48 months measured by the CBCL. However, we did not observe associations between neonatal brain organisation and developmental outcomes at 18 months. This finding is in line with previous neonatal fMRI studies that have reported inconsistent relationships between neonatal functional organisation and outcomes during toddlerhood. One possible explanation is that behavioural phenotypes are still emerging during the first few years, such that assessments during infancy may not fully capture individual differences in underlying brain organisation. In contrast, assessments during childhood may more reliably reflect these underlying differences. An alternative explanation is that “static” functional connectivity may not capture all aspects of early brain organisation. Recent work using the same dHCP cohort has shown that neonatal dynamic functional connectivity was associated with 18-month neurodevelopmental outcomes (França et al., 2024), suggesting that such dynamic measures may provide additional information beyond static connectivity alone.

This study has several limitations. First, follow-up data at 4-6 years were available for only 95 children. Although participants with and without follow-up data were broadly comparable, differences in PMA at scan suggest that the follow-up subset may be a self-selecting group that may not fully represent the whole sample. The limited sample size also makes the observed association with outcomes vulnerable to sample variability. We used the Schaefer atlas with corresponding Yeo-7 networks to ensure generalisability. While this atlas has been used in neonatal studies (King et al., 2023; Truzzi & Cusack, 2023; Yates et al., 2023; Yin et al., 2025), future studies may benefit from using neonate-specific atlases or precision mapping methods (Zhao et al., 2025). This is of particular relevance as the brain undergoes rapid development during the first years of life and thus adult parcellations may not accurately reflect the organisation of the neonatal brain. Second, functional integration and segregation were quantified using normalised global and local efficiency. While these graph-theoretical measures describe network organisation, they do not provide information about the specific within-network connectivity or interactions between functional networks underlying the observed effects.

In conclusion, we demonstrated that functional brain organisation undergoes rapid development during the first postnatal month and are disrupted by preterm birth. We further showed that neonatal functional organisation is associated with neurodevelopmental outcomes at 4-6 years, suggesting that individual differences in neonatal functional organisation are relevant to later development.

## Declaration of the use of AI

The authors declare the use of generative AI in the research process. According to the GAIDeT taxonomy (2025), code optimisation and proofreading task was delegated to GAI tools under full human supervision. The GAI tool used was GPT-5. Responsibility for the final manuscript lies entirely with the authors. GAI tools are not listed as authors and do not bear responsibility for the final outcomes.

## Data and Code Availability

The data used in this study were obtained from the Developing Human Connectome Project (dHCP) and are available through the dHCP data repository: https://data.developingconnectome.org.

All code and the 18-month follow-up data used in this study is available at https://github.com/QianwenChang/Neonatal_functional_organisation

The 4-6 year follow-up data from the CHiP project cannot be made publicly available due to ethical restrictions. Data will be shared upon reasonable request.

## Author Contributions

Q.C. and D.B. designed and performed research. Q.C. analysed data. S.F.M. pre-processed data. A.C., A.B., and B.G.E. acquired and processed follow-up data. C.N., and S.J.C, obtained funding for the follow-up study. Q.C. and D.B. wrote the manuscript. Q.C., S.F.M., A.C., A.B., B.G.E., C.N., T.A., G.M., S.J.C., and D.B. helped to interpret results, edited, and reviewed the manuscript. G.M., S.J.C., and D.B. supervised the study.

## Acknowledgement

The developing Human Connectome Project was supported by the European Research Council under the European Union’s Seventh Framework Programme (FP7/20072013)/ERC grant agreement no. 319456. Early childhood follow-up data acquisition was funded by the Medical Research Council UK [MR/V002465/1]. This work was also funded through philanthropic support from Heart of Racing LLC to the Brain Health Gen2020 programme at King’s College London, and by the National Institute for Health Research (NIHR) Biomedical Research Centre (BRC): Maudsley. The authors also acknowledge support from the Institute for Translational Neurodevelopment at King’s College London, the European Autism Interventions (EU-AIMS) trial and the EU AIMS-2-TRIALS, a European Innovative Medicines Initiative Joint Undertaking under Grant Agreements No. 115300 and 777394, the resources of which are composed of financial contributions from the European Union’s Seventh Framework Programme (Grant FP7/2007–2013) and Horizon 2020 from the European Federation of Pharmaceutical industries and Associations companies’ in-kind contributions. QC received support from KCL-CSC joint scholarship awarded by King’s College London and Chinese Scholarship Council. TA and GM received support from the Medical Research Council Centre for Neurodevelopmental Disorders, King’s College London [MR/N026063/1]. TA was supported by a Medical Research Council (MRC) Senior Clinical Fellowship [MR/Y009665/1]. The views expressed are those of the authors and not necessarily those of the funders, IHI-JU2, the NHS, the National Institute for Health Research, or the Department of Health and Social Care. The funders had no role in the design and conduct of the study; collection, management, analysis, and interpretation of the data; preparation, review, or approval of the manuscript; and decision to submit the manuscript for publication.

## Supplementary Material

**Supplementary Table S1.** Associations between functional network features at birth and neurodevelopmental outcomes at 4-6 years.

|  | Normalised global efficiency |  |  | Normalised local efficiency |  |  |
| --- | --- | --- | --- | --- | --- | --- |
|  | *r* | *p*<sub>UNC</sub> | *p*<sub>FDR</sub> | *r* | *p*<sub>UNC</sub> | *p*<sub>FDR</sub> |
| **SDQ total difficulties score** | **-0.306** | **0.003** | **0.035** | **-0.302** | **0.004** | **0.035** |
| Prosocial actions | 0.150 | 0.158 | 0.416 | 0.159 | 0.135 | 0.416 |
| Attention to others' feelings | -0.016 | 0.882 | 0.882 | -0.053 | 0.617 | 0.740 |
| Emotional contagion | -0.068 | 0.522 | 0.681 | -0.087 | 0.416 | 0.681 |
| SCQ total score | -0.071 | 0.507 | 0.681 | -0.095 | 0.372 | 0.681 |
| ADHD total score | -0.091 | 0.394 | 0.681 | -0.149 | 0.162 | 0.416 |
| Effortful control | 0.030 | 0.776 | 0.821 | 0.034 | 0.750 | 0.821 |
| Negative affectivity | -0.237 | 0.024 | 0.146 | -0.216 | 0.040 | 0.182 |
| Surgency | -0.067 | 0.529 | 0.681 | -0.083 | 0.438 | 0.681 |

**Note:** Spearman's partial correlations were calculated, controlling for corrected age at assessment, PMA at scan, head motion (FD outliers), sex, and IMD. *p*<sub>UNC</sub>, uncorrected *p*-value. *p*<sub>FDR</sub>, *p*-value adjusted for multiple comparisons using the Benjamini-Hochberg FDR method.

**Supplementary Table S2.** Demographic information of neonates with and without 18-month follow-up.

|  | With follow-up<br>(n = 303) | Without follow-up<br>(n = 91) | Statistic | p-value |
| --- | --- | --- | --- | --- |
| GA at birth [weeks] | 38.88 (3.24) | 38.38 (3.46) | 1.440 | 0.149 |
| PMA at scan [weeks] | 41.47 (1.80) | 41.31 (1.76) | 0.690 | 0.490 |
| Sex (male count, %) | 154 (50.83%) | 60 (65.93%) | 5.844 | 0.016* |
| Preterm birth [count, %] | 47 (15.51%) | 22 (24.18%) | 3.062 | 0.080 |
| IMD | 1454.44 (753.14) | 1223.93 (746.13) | 2.820 | 0.005** |
| % FD outliers | 5.25 (2.60) | 5.20 (2.71) | 0.153 | 0.879 |

**Note:** values are presented as mean (SD) if not indicated otherwise.
Chi-square test for sex and preterm birth, Mann-Whitney U test for other variables, \* $p < 0.05$ , \*\* $p < 0.01$ , \*\*\* $p < 0.001$ .

**Supplementary Table S3.** Demographic information of neonates with and without 4-6-year follow-up.

|  | With follow-up<br>(n = 95) | Without follow-up<br>(n = 299) | Statistic | p-value |
| --- | --- | --- | --- | --- |
| GA at birth [weeks] | 38.67 (3.38) | 38.80 (3.27) | 0.234 | 0.816 |
| PMA at scan [weeks] | 41.91 (1.66) | 41.28 (1.81) | 2.980 | 0.003** |
| Sex (male count, %) | 45 (47.37%) | 169 (56.52%) | 2.079 | 0.149 |
| Preterm birth [count, %] | 19 (20.00%) | 50 (16.72%) | 0.333 | 0.564 |
| IMD | 1443.22 (727.17) | 1388.96 (766.77) | 0.902 | 0.367 |
| % FD outliers | 4.82 (2.90) | 5.37 (2.52) | -1.790 | 0.074 |

**Note:** values are presented as mean (SD) if not indicated otherwise.
Chi-square test for sex and preterm birth, Mann-Whitney U test for other variables, \* $p < 0.05$ , \*\* $p < 0.01$ , \*\*\* $p < 0.001$ .

**Supplementary Table S4.** Demographic information of *term-born neonates* with and without 18-month follow-up.

|  | With follow-up<br>(n = 256) | Without follow-up<br>(n = 69) | Statistic | p-value |
| --- | --- | --- | --- | --- |
| GA at birth [weeks] | 40.06 (1.26) | 39.98 (1.25) | 0.407 | 0.684 |
| PMA at scan [weeks] | 41.61 (1.70) | 41.57 (1.54) | 0.295 | 0.769 |
| Sex (male count, %) | 129 (50.39%) | 45 (65.22%) | 4.226 | 0.040* |
| IMD | 1404.91 (738.88) | 1216.07 (732.10) | 2.130 | 0.033* |
| % FD outliers | 5.37 (2.60) | 5.39 (2.73) | -0.088 | 0.930 |

**Note:** values are presented as mean (SD) if not indicated otherwise.
Chi-square test for sex, Mann-Whitney U test for other variables, \* $p < 0.05$ , \*\* $p < 0.01$ , \*\*\* $p < 0.001$ .

**Supplementary Table S5.**
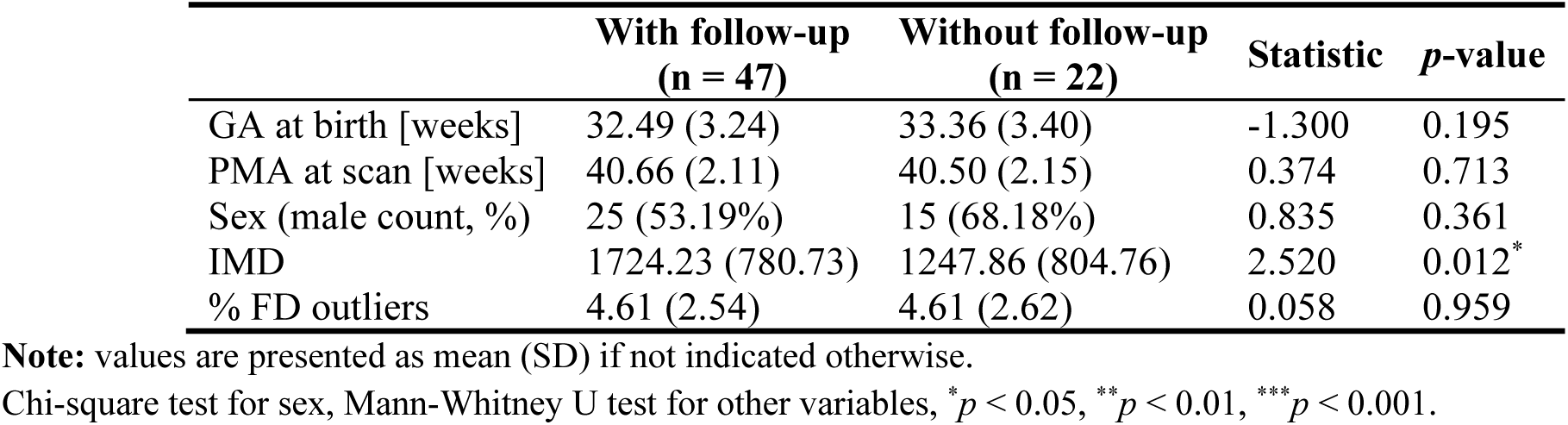
Demographic information of *preterm-born neonates* with and without 18-month follow-up.

|  | With follow-up<br>(n = 47) | Without follow-up<br>(n = 22) | Statistic | p-value |
| --- | --- | --- | --- | --- |
| GA at birth [weeks] | 32.49 (3.24) | 33.36 (3.40) | -1.300 | 0.195 |
| PMA at scan [weeks] | 40.66 (2.11) | 40.50 (2.15) | 0.374 | 0.713 |
| Sex (male count, %) | 25 (53.19%) | 15 (68.18%) | 0.835 | 0.361 |
| IMD | 1724.23 (780.73) | 1247.86 (804.76) | 2.520 | 0.012* |
| % FD outliers | 4.61 (2.54) | 4.61 (2.62) | 0.058 | 0.959 |

**Note:** values are presented as mean (SD) if not indicated otherwise.
Chi-square test for sex, Mann-Whitney U test for other variables, \* $p < 0.05$ , \*\* $p < 0.01$ , \*\*\* $p < 0.001$ .

**Supplementary Table S6.** Demographic information of *term-born neonates* with and without 4-6-year follow-up.

|  | With follow-up<br>(n = 76) | Without follow-up<br>(n = 249) | Statistic | p-value |
| --- | --- | --- | --- | --- |
| GA at birth [weeks] | 40.20 (1.21) | 39.99 (1.27) | 1.070 | 0.284 |
| PMA at scan [weeks] | 42.13 (1.56) | 41.44 (1.67) | 3.190 | 0.001** |
| Sex (male count, %) | 35 (46.05%) | 139 (55.82%) | 1.859 | 0.173 |
| IMD | 1382.86 (714.37) | 1360.47 (749.46) | 0.465 | 0.642 |
| % FD outliers | 4.93 (2.93) | 5.51 (2.51) | -1.600 | 0.111 |

**Note:** values are presented as mean (SD) if not indicated otherwise.
Chi-square test for sex, Mann-Whitney U test for other variables, \* $p < 0.05$ , \*\* $p < 0.01$ , \*\*\* $p < 0.001$ .

**Supplementary Table S7.** Demographic information of *preterm-born neonates* with and without 4-6-year follow-up.

|  | With follow-up<br>(n = 19) | Without follow-up<br>(n = 50) | Statistic | p-value |
| --- | --- | --- | --- | --- |
| GA at birth [weeks] | 32.56 (2.08) | 32.85 (3.67) | -1.020 | 0.310 |
| PMA at scan [weeks] | 41.01 (1.81) | 40.46 (2.21) | 0.961 | 0.340 |
| Sex (male count, %) | 10 (52.63%) | 30 (60.00%) | 0.079 | 0.779 |
| IMD | 1684.68 (747.04) | 1529.66 (841.08) | 0.927 | 0.357 |
| % FD outliers | 4.40 (2.81) | 4.69 (2.47) | -0.591 | 0.559 |

**Note:** values are presented as mean (SD) if not indicated otherwise.
Chi-square test for sex, Mann-Whitney U test for other variables, \* $p < 0.05$ , \*\* $p < 0.01$ , \*\*\* $p < 0.001$ .

## References

Achard, S., & Bullmore, E. (2007). Efficiency and Cost of Economical Brain Functional Networks. PLOS Computational Biology, 3(2), e17. 10.1371/journal.pcbi.0030017

Achenbach, T. M., & Ruffle, T. M. (2000). The Child Behavior Checklist and related forms for assessing behavioral/emotional problems and competencies. Pediatrics in Review, 21(8), 265–271. 10.1542/pir.21-8-265

Alcauter, S., Lin, W., Smith, J. K., Short, S. J., Goldman, B. D., Reznick, J. S., Gilmore, J. H., & Gao, W. (2014). Development of Thalamocortical Connectivity during Infancy and Its Cognitive Correlations. The Journal of Neuroscience, 34(27), 9067–9075. 10.1523/JNEUROSCI.0796-14.2014

Allison, C., Baron-Cohen, S., Wheelwright, S., Charman, T., Richler, J., Pasco, G., & Brayne, C. (2008). The Q-CHAT (Quantitative CHecklist for Autism in Toddlers): A normally distributed quantitative measure of autistic traits at 18-24 months of age: preliminary report. Journal of Autism and Developmental Disorders, 38(8), 1414–1425. 10.1007/s10803-007-0509-7

Avants, B. B., Tustison, N. J., Song, G., Cook, P. A., Klein, A., & Gee, J. C. (2011). A reproducible evaluation of ANTs similarity metric performance in brain image registration. NeuroImage, 54(3), 2033–2044. 10.1016/j.neuroimage.2010.09.025

Bashford-Largo, J., Nakua, H., Blair, R. J. R., Dominguez, A., Hatch, M., Blair, K. S., Dobbertin, M., Ameis, S., & Bajaj, S. (2024). A Shared Multivariate Brain-Behavior Relationship in a Transdiagnostic Sample of Adolescents. Biological Psychiatry: Cognitive Neuroscience and Neuroimaging, 9(4), 377–386. 10.1016/j.bpsc.2023.07.015

Batalle, D., Hughes, E. J., Zhang, H., Tournier, J.-D., Tusor, N., Aljabar, P., Wali, L., Alexander, D. C., Hajnal, J. V., Nosarti, C., Edwards, A. D., & Counsell, S. J. (2017). Early development of structural networks and the impact of prematurity on brain connectivity. NeuroImage, 149, 379–392. 10.1016/j.neuroimage.2017.01.065

Bayley, N. (2006). Bayley Scales of Infant and Toddler Development, Third Edition [Data set]. American Psychological Association (APA). 10.1037/t14978-000

Benjamini, Y., & Hochberg, Y. (1995). Controlling the False Discovery Rate: A Practical and Powerful Approach to Multiple Testing. Journal of the Royal Statistical Society: Series B (Methodological), 57(1), 289–300. 10.1111/j.2517-6161.1995.tb02031.x

Bouyssi-Kobar, M., De Asis-Cruz, J., Murnick, J., Chang, T., & Limperopoulos, C. (2019). Altered Functional Brain Network Integration, Segregation and Modularity in Very Preterm Infants at Term Equivalent Age. The Journal of Pediatrics, 213, 13–21.e1. 10.1016/j.jpeds.2019.06.030

Cao, M., Huang, H., & He, Y. (2017). Developmental Connectomics from Infancy through Early Childhood. Trends in Neurosciences, 40(8), 494–506. 10.1016/j.tins.2017.06.003

Chen, H., Huang, L., Yang, D., Ye, Q., Guo, M., Qin, R., Luo, C., Li, M., Ye, L., Zhang, B., & Xu, Y. (2019). Nodal Global Efficiency in Front-Parietal Lobe Mediated Periventricular White Matter Hyperintensity (PWMH)-Related Cognitive Impairment. Frontiers in Aging Neuroscience, 11, 347. 10.3389/fnagi.2019.00347

Chew, A. T. M., Bonthrone, A. F., Alford, A., Kelly, C., Pushparajah, K., Egloff, A., Hajnal, J. V., Simpson, J., Rutherford, M., Edwards, A. D., Nosarti, C., & Counsell, S. J. (2024). Executive Function in Preschool Children with Congenital Heart Disease and Controls: The Role of a Cognitively Stimulating Home Environment. The Journal of Pediatrics, 267, 113897. 10.1016/j.jpeds.2023.113897

Chew, A. T. M., Bonthrone, A. F., van der Meijden, M. E. M., Sun, Z., Pushparajah, K., Simpson, J., Edwards, A. D., Counsell, S. J., & Nosarti, C. (2026). Behavioral outcomes of preschool children with congenital heart disease and controls. Frontiers in Pediatrics, 14. 10.3389/fped.2026.1725994

Cohen, J. R., & D’Esposito, M. (2016). The Segregation and Integration of Distinct Brain Networks and Their Relationship to Cognition. Journal of Neuroscience, 36(48), 12083–12094. 10.1523/JNEUROSCI.2965-15.2016

Cole, M. W., Repovs, G., & Anticevic, A. (2014). The frontoparietal control system: A central role in mental health. *The Neuroscientist : A Review Journal Bringing Neurobiology*, Neurology and Psychiatry, 20(6), 652–664. 10.1177/1073858414525995

Derman, D., & Ferradal, S. L. (2025). Tracking functional brain networks in preterm and term infants using precision mapping. Developmental Cognitive Neuroscience, 76, 101629. 10.1016/j.dcn.2025.101629

Diedrichsen, J. (2006). A spatially unbiased atlas template of the human cerebellum. NeuroImage, 33(1), 127–138. 10.1016/j.neuroimage.2006.05.056

Disselhoff, V., Jakab, A., Latal, B., Schnider, B., Wehrle, F. M., & Hagmann, C. F. (2025). Inhibition abilities and functional brain connectivity in school-aged term-born and preterm-born children. Pediatric Research, 97(1), 315–324. 10.1038/s41390-024-03241-0

Edwards, A. D., Rueckert, D., Smith, S. M., Abo Seada, S., Alansary, A., Almalbis, J., Allsop, J., Andersson, J., Arichi, T., Arulkumaran, S., Bastiani, M., Batalle, D., Baxter, L., Bozek, J., Braithwaite, E., Brandon, J., Carney, O., Chew, A., Christiaens, D., … Hajnal, J. V. (2022). The Developing Human Connectome Project Neonatal Data Release. Frontiers in Neuroscience, 16, 886772. 10.3389/fnins.2022.886772

Eyre, M., Fitzgibbon, S. P., Ciarrusta, J., Cordero-Grande, L., Price, A. N., Poppe, T., Schuh, A., Hughes, E., O’Keeffe, C., Brandon, J., Cromb, D., Vecchiato, K., Andersson, J., Duff, E. P., Counsell, S. J., Smith, S. M., Rueckert, D., Hajnal, J. V., Arichi, T., … Edwards, A. D. (2021). The Developing Human Connectome Project: Typical and disrupted perinatal functional connectivity. Brain, 144(7), 2199–2213. 10.1093/brain/awab118

Fang, T., Wang, R., Liu, W., Zhang, Y., Guo, Y., Hu, Y., Zhao, X., Chen, Y., Fan, Q., & Ming, D. (2025). Edge participation coefficient unveiling the developmental dynamics of neonatal functional connectome. Communications Biology, 8(1), 1463. 10.1038/s42003-025-08873-4

Fenn-Moltu, S., Fitzgibbon, S. P., Ciarrusta, J., Eyre, M., Cordero-Grande, L., Chew, A., Falconer, S., Gale-Grant, O., Harper, N., Dimitrova, R., Vecchiato, K., Fenchel, D., Javed, A., Earl, M., Price, A. N., Hughes, E., Duff, E. P., O’Muircheartaigh, J., Nosarti, C., … Batalle, D. (2023). Development of neonatal brain functional centrality and alterations associated with preterm birth. Cerebral Cortex, 33(9), 5585–5596. 10.1093/cercor/bhac444

Fitzgibbon, S. P., Harrison, S. J., Jenkinson, M., Baxter, L., Robinson, E. C., Bastiani, M., Bozek, J., Karolis, V., Cordero Grande, L., Price, A. N., Hughes, E., Makropoulos, A., Passerat-Palmbach, J., Schuh, A., Gao, J., Farahibozorg, S.-R., O’Muircheartaigh, J., Ciarrusta, J., O’Keeffe, C., … Andersson, J. (2020). The developing Human Connectome Project (dHCP) automated resting-state functional processing framework for newborn infants. NeuroImage, 223, 117303. 10.1016/j.neuroimage.2020.117303

França, L. G. S., Ciarrusta, J., Gale-Grant, O., Fenn-Moltu, S., Fitzgibbon, S., Chew, A., Falconer, S., Dimitrova, R., Cordero-Grande, L., Price, A. N., Hughes, E., O’Muircheartaigh, J., Duff, E., Tuulari, J. J., Deco, G., Counsell, S. J., Hajnal, J. V., Nosarti, C., Arichi, T., … Batalle, D. (2024). Neonatal brain dynamic functional connectivity in term and preterm infants and its association with early childhood neurodevelopment. Nature Communications, 15(1), 16. 10.1038/s41467-023-44050-z

Gao, W. (2025). A hierarchical model of early brain functional network development. Trends in Cognitive Sciences. 10.1016/j.tics.2025.04.001

Gao, W., Alcauter, S., Smith, J. K., Gilmore, J. H., & Lin, W. (2015). Development of human brain cortical network architecture during infancy. Brain Structure and Function, 220(2), 1173– 1186. 10.1007/s00429-014-0710-3

Gao, W., Gilmore, J. H., Giovanello, K. S., Smith, J. K., Shen, D., Zhu, H., & Lin, W. (2011). Temporal and Spatial Evolution of Brain Network Topology during the First Two Years of Life. PLOS ONE, 6(9), e25278. 10.1371/journal.pone.0025278

Gilmore, J. H., Knickmeyer, R. C., & Gao, W. (2018). Imaging structural and functional brain development in early childhood. Nature Reviews Neuroscience, 19(3), 123–137. 10.1038/nrn.2018.1

Goodman, A., & Goodman, R. (2009). Strengths and Difficulties Questionnaire as a Dimensional Measure of Child Mental Health. Journal of the American Academy of Child & Adolescent Psychiatry, 48(4), 400–403. 10.1097/CHI.0b013e3181985068

Goodman, R. (1997). The Strengths and Difficulties Questionnaire: A Research Note. Journal of Child Psychology and Psychiatry, 38(5), 581–586. 10.1111/j.1469-7610.1997.tb01545.x

Goodman, R. (2001). Psychometric Properties of the Strengths and Difficulties Questionnaire. Journal of the American Academy of Child & Adolescent Psychiatry, 40(11), 1337–1345. 10.1097/00004583-200111000-00015

Gustavson, D. E., Borriello, G. A., Karhadkar, M. A., Rhee, S. H., Corley, R. P., Rhea, S.-A., DiLalla, L. F., Wadsworth, S. J., Friedman, N. P., & Reynolds, C. A. (2025). Stability of general cognitive ability from infancy to adulthood: A combined twin and genomic investigation. Proceedings of the National Academy of Sciences, 122(21), e2426531122. 10.1073/pnas.2426531122

Hadaya, L., Váša, F., Dimitrakopoulou, K., Saqi, M., Shergill, S. S., Edwards, A. D., Batalle, D., Leech, R., & Nosarti, C. (2025). Exploring functional connectivity in clinical and data-driven groups of preterm and term adults. Brain Communications, 7(2), fcaf074. 10.1093/braincomms/fcaf074

Hughes, E. J., Winchman, T., Padormo, F., Teixeira, R., Wurie, J., Sharma, M., Fox, M., Hutter, J., Cordero-Grande, L., Price, A. N., Allsop, J., Bueno-Conde, J., Tusor, N., Arichi, T., Edwards, A. D., Rutherford, M. A., Counsell, S. J., & Hajnal, J. V. (2017). A dedicated neonatal brain imaging system. Magnetic Resonance in Medicine, 78(2), 794–804. 10.1002/mrm.26462

Karolis, V. R., Froudist-Walsh, S., Brittain, P. J., Kroll, J., Ball, G., Edwards, A. D., Dell’Acqua, F., Williams, S. C., Murray, R. M., & Nosarti, C. (2016). Reinforcement of the Brain’s Rich-Club Architecture Following Early Neurodevelopmental Disruption Caused by Very Preterm Birth. Cerebral Cortex, 26(3), 1322–1335. 10.1093/cercor/bhv305

Keunen, K., Counsell, S. J., & Benders, M. J. N. L. (2017). The emergence of functional architecture during early brain development. NeuroImage, 160, 2–14. 10.1016/j.neuroimage.2017.01.047

King, G., Truzzi, A., & Cusack, R. (2023). The confound of head position in within-session connectome fingerprinting in infants. NeuroImage, 265, 119808. 10.1016/j.neuroimage.2022.119808

Li, Q., Xia, M., Zeng, D., Xu, Y., Sun, L., Liang, X., Xu, Z., Zhao, T., Liao, X., Yuan, H., Liu, Y., Huo, R., Li, S., & He, Y. (2024). Development of segregation and integration of functional connectomes during the first 1,000 days. Cell Reports, 43(5). 10.1016/j.celrep.2024.114168

López-Guerrero, N., & Alcauter, S. (2025). Developmental Trajectories and Differences in Functional Brain Network Properties of Preterm and At-Term Neonates. Human Brain Mapping, 46(1), e70126. 10.1002/hbm.70126

Lyu, W., Thung, K.-H., Huynh, K. M., Wang, L., Lin, W., Ahmad, S., & Yap, P.-T. (2025). Functional development of the human cerebellum from birth to age five. Nature Communications, 16(1), 6350. 10.1038/s41467-025-61465-y

Marek, S., & Dosenbach, N. U. F. (2018). The frontoparietal network: Function, electrophysiology, and importance of individual precision mapping. Dialogues in Clinical Neuroscience, 20(2), 133–140. 10.31887/DCNS.2018.20.2/smarek

McGoey, K. E., DuPaul, G. J., Haley, E., & Shelton, T. L. (2007). Parent and Teacher Ratings of Attention-Deficit/Hyperactivity Disorder in Preschool: The ADHD Rating Scale-IV Preschool Version. Journal of Psychopathology and Behavioral Assessment, 29(4), 269–276. 10.1007/s10862-007-9048-y

Michael, C., Mitchell, M. E., Cascone, A. D., Fogleman, N. D., Rosch, K. S., Cutts, S. A., Pekar, J. J., Sporns, O., Mostofsky, S. H., & Cohen, J. R. (2024). Reconfiguration of Functional Brain Network Organization and Dynamics With Changing Cognitive Demands in Children With Attention-Deficit/Hyperactivity Disorder. Biological Psychiatry: Cognitive Neuroscience and Neuroimaging. 10.1016/j.bpsc.2024.11.006

Oldham, S., Mansour L., S., & Ball, G. (2025). Perinatal development of structural thalamocortical connectivity. Imaging Neuroscience, 3, imag_a_00418. 10.1162/imag_a_00418

Onofrj, V., Chiarelli, A. M., Wise, R., Colosimo, C., & Caulo, M. (2022). Interaction of the salience network, ventral attention network, dorsal attention network and default mode network in neonates and early development of the bottom-up attention system. Brain Structure & Function, 227(5), 1843–1856. 10.1007/s00429-022-02477-y

Park, S., Haak, K. V., Oldham, S., Cho, H., Byeon, K., Park, B., Thomson, P., Chen, H., Gao, W., Xu, T., Valk, S., Milham, M. P., Bernhardt, B., Di Martino, A., & Hong, S.-J. (2024). A shifting role of thalamocortical connectivity in the emergence of cortical functional organization. Nature Neuroscience, 27(8), 1609–1619. 10.1038/s41593-024-01679-3

Putnam, S. P., Helbig, A. L., Gartstein, M. A., Rothbart, M. K., & Leerkes, E. (2014). Development and assessment of short and very short forms of the infant behavior questionnaire-revised. Journal of Personality Assessment, 96(4), 445–458. 10.1080/00223891.2013.841171

Rieffe, C., Ketelaar, L., & Wiefferink, C. H. (2010). Assessing empathy in young children: Construction and validation of an Empathy Questionnaire (EmQue). Personality and Individual Differences, 49(5), 362–367. 10.1016/j.paid.2010.03.046

Rubinov, M., & Sporns, O. (2010). Complex network measures of brain connectivity: Uses and interpretations. NeuroImage, 52(3), 1059–1069. 10.1016/j.neuroimage.2009.10.003

Rudie, J. D., Brown, J. A., Beck-Pancer, D., Hernandez, L. M., Dennis, E. L., Thompson, P. M., Bookheimer, S. Y., & Dapretto, M. (2013). Altered functional and structural brain network organization in autism. NeuroImage: Clinical, 2, 79–94. 10.1016/j.nicl.2012.11.006

Sa de Almeida, J., Meskaldji, D.-E., Loukas, S., Lordier, L., Gui, L., Lazeyras, F., & Hüppi, P. S. (2021). Preterm birth leads to impaired rich-club organization and fronto-paralimbic/limbic structural connectivity in newborns. NeuroImage, 225, 117440. 10.1016/j.neuroimage.2020.117440

Schaefer, A., Kong, R., Gordon, E. M., Laumann, T. O., Zuo, X.-N., Holmes, A. J., Eickhoff, S. B., & Yeo, B. T. T. (2018). Local-Global Parcellation of the Human Cerebral Cortex from Intrinsic Functional Connectivity MRI. Cerebral Cortex, 28(9), 3095–3114. 10.1093/cercor/bhx179

Schafer, R. J., Lacadie, C., Vohr, B., Kesler, S. R., Katz, K. H., Schneider, K. C., Pugh, K. R., Makuch, Robert. W., Reiss, A. L., Constable, R. T., & Ment, L. R. (2009). Alterations in functional connectivity for language in prematurely born adolescents. Brain, 132(3), 661– 670. 10.1093/brain/awn353

Scheinost, D., Chang, J., Brennan-Wydra, E., Lacadie, C., Constable, R. T., Chawarska, K., & Ment, L. R. (2024). Developmental trajectories of the default mode, frontoparietal, and salience networks from the third trimester through the newborn period. Imaging Neuroscience, 2, 1– 16. 10.1162/imag_a_00201

Schuh, A., Makropoulos, A., Robinson, E. C., Cordero-Grande, L., Hughes, E., Hutter, J., Price, A. N., Murgasova, M., Teixeira, R. P. A. G., Tusor, N., Steinweg, J. K., Victor, S., Rutherford, M. A., Hajnal, J. V., Edwards, A. D., & Rueckert, D. (2018). Unbiased construction of a temporally consistent morphological atlas of neonatal brain development (p. 251512). bioRxiv. 10.1101/251512

Smyser, C. D., Inder, T. E., Shimony, J. S., Hill, J. E., Degnan, A. J., Snyder, A. Z., & Neil, J. J. (2010). Longitudinal analysis of neural network development in preterm infants. Cerebral Cortex, 20(12), 2852–2862. 10.1093/cercor/bhq035

Sporns, O. (2013). Network attributes for segregation and integration in the human brain. *Current Opinion in Neurobiology*, Macrocircuits, 23(2), 162–171. 10.1016/j.conb.2012.11.015

Tikoo, S., Hernandez-Castillo, C. R., Chen, H., Stephens, R., Cornea, E., Gilmore, J. H., & Gao, W. (2025). The Evolving Cerebellar and Cerebello-cortical Functional Connectivity Architecture during Infancy. The Journal of Neuroscience: The Official Journal of the Society for Neuroscience, 45(11), e1209242025. 10.1523/JNEUROSCI.1209-24.2025

Toulmin, H., Beckmann, C. F., O’Muircheartaigh, J., Ball, G., Nongena, P., Makropoulos, A., Ederies, A., Counsell, S. J., Kennea, N., Arichi, T., Tusor, N., Rutherford, M. A., Azzopardi, D., Gonzalez-Cinca, N., Hajnal, J. V., & Edwards, A. D. (2015). Specialization and integration of functional thalamocortical connectivity in the human infant. Proceedings of the National Academy of Sciences, 112(20), 6485–6490. 10.1073/pnas.1422638112

Truzzi, A., & Cusack, R. (2023). The development of intrinsic timescales: A comparison between the neonate and adult brain. NeuroImage, 275, 120155. 10.1016/j.neuroimage.2023.120155

Vanes, L., Fenn-Moltu, S., Hadaya, L., Fitzgibbon, S., Cordero-Grande, L., Price, A., Chew, A., Falconer, S., Arichi, T., Counsell, S. J., Hajnal, J. V., Batalle, D., Edwards, A. D., & Nosarti, C. (2023). Longitudinal neonatal brain development and socio-demographic correlates of infant outcomes following preterm birth. Developmental Cognitive Neuroscience, 61, 101250. 10.1016/j.dcn.2023.101250

Wang, R., Liu, M., Cheng, X., Wu, Y., Hildebrandt, A., & Zhou, C. (2021). Segregation, integration, and balance of large-scale resting brain networks configure different cognitive abilities. Proceedings of the National Academy of Sciences, 118(23), e2022288118. 10.1073/pnas.2022288118

Weber, C. F., Kebets, V., Benkarim, O., Lariviere, S., Wang, Y., Ngo, A., Jiang, H., Chai, X., Park, B., Milham, M. P., Di Martino, A., Valk, S., Hong, S.-J., & Bernhardt, B. C. (2024). Contracted functional connectivity profiles in autism. Molecular Autism, 15(1), 38. 10.1186/s13229-024-00616-2

Wee, C.-Y., Tuan, T. A., Broekman, B. F. P., Ong, M. Y., Chong, Y.-S., Kwek, K., Shek, L. P.-C., Saw, S.-M., Gluckman, P. D., Fortier, M. V., Meaney, M. J., & Qiu, A. (2017). Neonatal neural networks predict children behavioral profiles later in life. Human Brain Mapping, 38(3), 1362–1373. 10.1002/hbm.23459

Xie, C., Xiang, S., Shen, C., Peng, X., Kang, J., Li, Y., Cheng, W., He, S., Bobou, M., Broulidakis, M. J., van Noort, B. M., Zhang, Z., Robinson, L., Vaidya, N., Winterer, J., Zhang, Y., King, S., Banaschewski, T., Barker, G. J., … Feng, J. (2023). A shared neural basis underlying psychiatric comorbidity. Nature Medicine, 29(5), 1232–1242. 10.1038/s41591-023-02317-4

Xu, Y., Zhao, T., Jeon, T., Ouyang, M., Zhu, T., Chalak, L., Rollins, N., & Huang, H. (2026). Structural and Functional Coupling in Neonates and its Prediction for Neurocognitive Outcomes. Journal of Pacific Rim Psychology, 20, 18344909251414889. 10.1177/18344909251414889

Yates, T. S., Ellis, C. T., & Turk-Browne, N. B. (2023). Functional networks in the infant brain during sleep and wake states. Cerebral Cortex, 33(21), 10820–10835. 10.1093/cercor/bhad327

Yeo, B. T. T., Krienen, F. M., Sepulcre, J., Sabuncu, M. R., Lashkari, D., Hollinshead, M., Roffman, J. L., Smoller, J. W., Zöllei, L., Polimeni, J. R., Fischl, B., Liu, H., & Buckner, R. L. (2011). The organization of the human cerebral cortex estimated by intrinsic functional connectivity. Journal of Neurophysiology, 106(3), 1125–1165. 10.1152/jn.00338.2011

Yin, W., Li, T., Wu, Z., Hung, S.-C., Hu, D., Gui, Y., Cho, S., Sun, Y., Woodburn, M. A., Wang, L., Li, G., Piven, J., Elison, J. T., Wu, C. W., Zhu, H., Cohen, J. R., & Lin, W. (2025). Charting brain functional development from birth to 6 years of age. Nature Human Behaviour, 1–14. 10.1038/s41562-025-02160-2

Zhao, J., Xu, Y., Cui, Z., Li, H., Sun, L., Liang, X., Han, M., Zeng, Z., Li, Q., Zhao, T., & He, Y. (2025). Emergence of Individualized Functional Topography in the Neonatal Brain (p. 2025.11.18.689151). bioRxiv. 10.1101/2025.11.18.689151

Zhao, T., Xu, Y., & He, Y. (2019). Graph theoretical modeling of baby brain networks. NeuroImage, 185, 711–727. 10.1016/j.neuroimage.2018.06.038

